# Exploring the effects of high concentration of glucose on soil microbial community: An experimental study using the soil of North Bangalore

**DOI:** 10.1101/017061

**Authors:** Anshuman Swain

## Abstract

Substrate Induced Respiration (SIR) is a standard method to study microbial biomass in soil. It is observed that the soil microbial CO_2_ respiration goes up with the glucose concentration till a certain concentration, and afterwards decreases and stabilizes. There are two possible mechanisms via which this can happen: increased osmotic pressure can kill off a group of microbial population or the Crabtree effect takes over the population. An experiment was designed using the SIR; to find the reason for the same and prove or disprove one of these hypothesises.

## Introduction

The technique of substrate induced respiration (SIR) has been designed to evaluate the degree of respiration in the soil due the employment of a substrate such as glucose, glutamic acid, mannitol and amino acids. The biological respiration reactions of the organisms present in the soil and their by-products, due to substrate addition, such as production of CO_2_ and/or consumption of O_2_, are used in this method as a measure of calculating the microbial activities in the soil. Anderson and Domsch came up with this process in 1978 to provide a swift valuation of the live microbial biomass in soils. The substrates that can be utilized are not limited to the aforesaid ones only; the choice of the substrate that is used should reflect the organisms that are in target and soil type that is being tested.

When this method of SIR is employed on a soil of North Bangalore, it was found that the total CO_2_ release due to respiration increased with the increase in glucose concentration but after it reached 0.4g/10 g of soil sample, it started to decrease fast and stabilized at a lower value. This means that the net respiration processes have decreased which can be a result of, a decrease in the total number of organisms (the rate of respiration remaining the same), a decrease in the rate of respiration (the number of organisms remaining the same) or a mixture of both the above reasons.

The first one will be due to high osmotic pressure (of glucose), which can cause the cells to die as they will not be able to withstand such a highly hypotonic solution. The second one will be due to Crabtree effect, in which many of the organisms generate the ATP using glycolysis only without going on to the Kreb’s cycle and hence producing negligible CO_2_.

## Method

Substrate induced respiration is used on the North Bangalore soil and the results of the carbon dioxide released versus the initial concentration of glucose is plotted and the graph is shown in figure 1.

**Figure 1.**
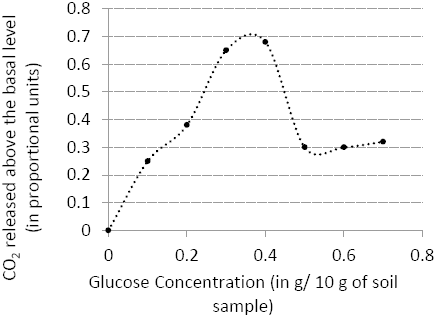

The reason for the decrease in the CO_2_ emission after the concentration of 0.4g/ 10g of soil sample was reached, plausibly could be either the extreme osmotic pressure of the saturation of glucose or the Crabtree effect, To test which of the above two may be the reason of the decrease in CO_2_ emission, an experiment was designed through which it could be deduced whether or not in the higher concentration the organisms die or not.

The basic idea behind the experiment is to incubate the microbes in high glucose concentration and measure the CO_2_ emissions using SIR and then decrease the glucose concentration in the same sample & incubate. Then it is compared with a control sample with the concentration, to which it was reduced later, which is also made to undergo SIR.

## Experiment

The protocol was followed throughout the experiment involving the loamy soil collected from a site in Northern Bangalore (Karnataka, India). It had pH 8.34 and a water holding capacity of 4 mL/10 g of soil. It was freshly collected and used, thus did not require any special incubation or other treatment. In SIR, the soil samples were measured at 10gm and put in a conical flask, then glucose of a certain concentration was added to it at 40% water holding capacity, and the mouth of the flask was corked and there was a tube through the cork connecting to another similar flask tilted horizontally at a higher elevation that containing 10 mL NaOH solution of a certain concentration. The CO_2_ that will be released by the microbes in the flask will be absorbed by the NaOH solution in the other flask. Then the amount of CO_2_ released and that has been absorbed by the NaOH solution, is determined by titration against HCl of known concentration after addition BaCl_2_ or Ba(NO_3_)_2_ in order to precipitate the carbonate, and using phenolphthalein as the indicator, so as to determine the amount of NaOH left and hence the amount of NaOH used up in reaction against CO_2_.

Hourly CO_2_ emissions are checked using SIR for a 0.3 g/10g soil control concentration and a 0.6g/10 g test concentration for five hours and then after five hours, fresh 10 g of soil wetted with 1.6mL distilled water is put in the test flasks containing the soil with higher glucose concentration. The experiment, as seen in figure 2 was repeated for 3 times and values were noted.

**Figure 2.**
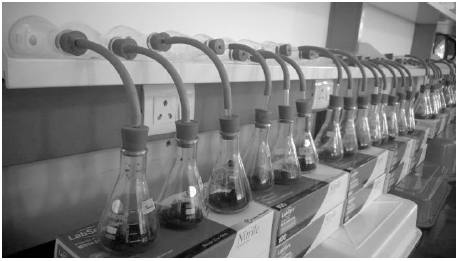
The Experimental Setup.

## Results and Discussion

All the values were statistically analysed using Matlab and the data was plotted as shown in figure 3.

**Figure 3.**
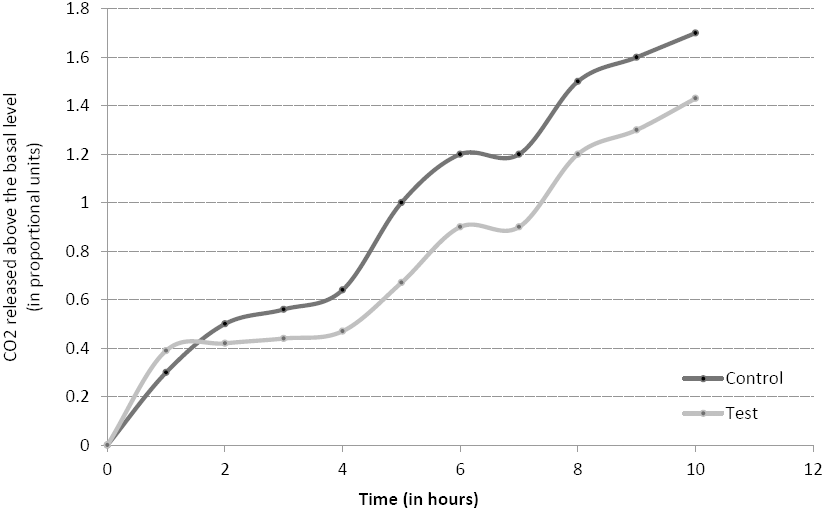
Comparative time varied graph.

This graph actually disproves the osmotic pressure hypothesis and proves the Crabtree effect. As seen in the beginning, the rate of CO_2_ emission is higher in case of higher glucose concentration (test sample) but the total amount soon becomes less than the control sample, this is what was observed in figure 1, and this continues till 5 hrs, after which both the concentrations are made equal, and had the increased concentration killed the microbes, there would have been lesser carbon dioxide emissions after 5 hrs also, but as it is seen in the graph the amount of CO_2_ emitted after 5 hrs till the end of the experiment is same for both the cases, and hence the total number of organisms is nearly the same. The number of organisms in both the sample is same despite the fact that more soil was added, is that, after 5 hours, there would be the second generation microbes, which will be *2^n+1^* if the initial number is *n* and moreover *n* is quite large, hence *2^n+1^* is much greater than *n*. So when the new soil is added after 5 hours, the total number of microbes in the control sample is *2^n+1^* whereas that in the test sample is *2^n+1^+n*, which is approximately equal to *2^n+1^*.

So, from the results of figure 1, we can see that 57.14% of the total microbe population were affected by the Crabtree effect, as had there been no such effect; the total CO_2_ emission would have remained constant, even though the concentration would have increased and that would have signified the concentration of maximal glucose consumption and the decrease is due to Crabtree effect.

In addition, as the rate of CO_2_ emission is same for both cases after 5 hours, there is no observable decrease in the number of organisms, which would have been due to the deaths by high osmotic pressure.

## Conclusion

From the designed experiment, it can be safely concluded that at high glucose concentrations, the microbes are affected by the Crabtree effect, in which they use the carbon nutrient source anaerobically to produce ATP and thus don’t produce CO_2_. So, SIR cannot be satisfactorily be used for soils with high glucose dosage as it will give undermining results.

## Acknowledgements

We are grateful to Dr Sumanta Bagchi for giving us an opportunity to work in his lab and also to Mr Manjunatha H C, for his great help in setting up the experiment and to other people in his lab for their sincere cooperation.

